# Identification and validation of a ferroptosis-related genes based prognostic signature for prostate cancer

**DOI:** 10.1101/2020.10.17.343897

**Authors:** Huan Liu, Lei Gao, Jie Li, Tingshuai Zhai, Tiancheng Xie, Yunfei Xu

## Abstract

Ferroptosis, an iron-dependent form of selective cell death, involves in the development of many cancers. However, systematic analysis of ferroptosis related genes (FRGs) in prostate cancer (PCa) remains to be clarified. In our research, we collected the mRNA expression profiles and clinical information of PCa patients from TCGA and MSKCC databases. The univariate, LASSO and multivariate Cox regression method were performed to construct prognostic signature in TCGA cohort. Seven FRGs, AKR1C3, ALOXE3, ATP5MC3, CARS1, MT1G, PTGS2, TFRC, were included to establish the risk model, which was validated in MSKCC dataset. Subsequently, we found that high risk group was strongly correlated with copy number alteration load, tumor burden mutation, immune cell infiltration, mRNAsi, immuetherapy and bicalutamide response. Finally, it was identified that overexpression of TFRC could induce proliferation and invasion in PCa cell lines *in vitro.* These results demonstrated that this risk model based on recurrence free survival (RFS) could accurately predict prognosis in PCa patients, suggesting that FRGs are promising prognostic biomarkers and drug target genes for PCa patients.

## 1. Introduction

Prostate cancer (PCa) is one of the most common male malignancies among the world, which caused the second cancer-related deaths across western countries [1]. Since 1990s, prostate-specific antigen (PSA) has been used as the standard test for PCa detection. However, there were no significant differences in mortality among PSA-screened patients comparing with those without screening [2–4]. PSA is also regarded as a significant sign for prognosis of PCa. For those patients accepted radical prostatectomy and radiotherapy, 27%-53% of them experienced the return of PSA, which was defined as biochemical recurrence (BCR) [5]. The BCR will lead to the development of advanced castration-resistant prostate cancer (CRPC) stage and resulted in an increasement of risk in distant metastases, prostate cancer-specific mortality and overall mortality [6, 7]. Hence, it is of great significance to identify novel prognostic biomarkers for PCa.

Ferroptosis, a newly identified form of regulated cell death (RCD) characterized by iron accumulation and lipid peroxidation, is distinct from other forms of RCD (necroptosis, apoptosis, or autophagic cell death) [8]. Recently, emerging evidence suggested that ferroptosis is related with cancer initiation, progression or drug sensitivity [9–11]. Application of ferroptosis inducers could help overcoming drug resistance and inhibiting cancer progression or metastasis. For example, Tang et al reported that knockdown of metallothionein-1G (MT-1G) enhances the sensitivity of sorafenib in hepatocellular carcinoma through promotion of ferroptosis [10]. Shi’s group indicated that cysteine dioxygenase 1 (CDO1) suppression increased cellular glutathione (GSH) levels, inhibited reactive oxygen species (ROS) generation, and decreased lipid peroxidation in erastin-treated gastric cancer cells [12].

The role of ferroptosis in PCa has drawn attention in recent years. Butler et al’s study revealed that knockdown of DECR1 in PCa cells inhibits tumor cells proliferation and migration via accumulating cellular polyunsaturated fatty acids (PUFAs), enhancing mitochondrial oxidative stress and lipid peroxidation, and promoting ferroptosis [13]. In Pan’s research, the clarified pannexin 2 (PANX2) as a new marker which regulates ferroptosis via Nrf2 signaling pathway and accelerates cancer cell proliferation, invasion in PCa [14]. Though preliminary evidence has identified several markers correlated with ferroptosis in PCa, the association between other ferroptosis-related genes (FRGs) and PCa patient prognosis still remains largely unknown.

In our research, we firstly collected the mRNA expression profiles of 40 ferroptosis-related genes and clinical data of PCa patients from the TCGA database. Then we evaluated their differential expression in different risk PCa samples and investigated the enriched pathways and biological roles. Moreover, we chose the hub gene transferrin receptor (TFRC) for further experiment *in vitro.* As a result, we constructed a FRGs-based prognostic model in PCa of TCGA dataset and validated it in other dataset, and verified the TFRC gene function, which might strengthen our understanding of PCa.

## 2. Materials and methods

### 2.1. Data collection

The RNA-seq (FPKM value) data of 499 PCa and 52 normal prostate tissues with related clinical data were collected from the TCGA website (https://portal.gdc.cancer.gov/repository). As validation dataset, the MSKCC data were enrolled as the validation cohort downloaded from GEO dataset (https://www.ncbi.nlm.nih.gov/geo/query/), which included integrated genomic profiling of 218 prostate tumors. All datasets used in this study could be reached to the public.

### 2.2. Gene Signature Building

40 genes were included in this study which were verified in the research of Stockwell et al [15] and are provided in Supplementary Table S1. Univariate Cox was performed to select the RFS related genes, and genes with P values less than 0.05 were retained. Then LASSO method was applied to minimize the risk of overfitting. Finally, multiple stepwise Cox regression was utilized to establish the risk model. The risk score of hub genes was established as (exprgene1 × coefficientgene1) + (exprgene2 × coeficientgene2) + ⋯ + (exprgene7 × coefficientgene7).

### 2.3. Clustering, genetic alterations, functional enrichment analysis

Consensus clustering based on ferroptosis genes was identified using “ Consensus ClusterPlus” package [16]. The “clusterProfiler” R package [17] was applied to perform GO and KEGG analyses based on the different expression genes (DEGs) (|log2FC| ≥ 1, FDR < 0.05) between different risk groups. The infiltrating score of 28 immune cells were calculated with ssGSEA in the “GSVA” R package [18]. The annotated gene set file is provided in Supplementary Table S2. The genetic alterations of selective genes in TCGA patients were acquired from the cBioPortal [19, 20]. The relationships between mRNA expression level and gleason score, lymph nodal metastasis, methylation levels were acquired from UALCAN[21].

### 2.4. Copy number variation (CNV) load, tumor mutation burden (TMB), neoantigens, tumor stemness and clonal score

Based on the median value of risk score, the TCGA patients were divided into two risk group. Then the CNV load at the focal and arm levels were calculated based on the GISTIC_2.0 results, which were freely available from Broad Firehose (https://gdac.broadinstitute.org/). The tumor mutation burden (TMB) of each patient was calculated as the total number of non-synonymous mutations per megabase. Tumor neoantigens, which could be recognized by neoantigen-spcific T cell receptors (TCRs) and may play critical roles in T-cell-mediated antitumor immune response, were analyzed based on the results from TCIA (https://tcia.at/). Clonality is the critical character of tumors. Then we also analyzed the clonality of PCa patients from TCIA dataset. Moreover, considering the tumor stemness is also one of the basic traits of tumor, then mRNAsi data was analyzed based on the report of Robertson [22].

### 2.5. Prediction of immunotherapy and chemotherapy responses

In order to assess the clinical response to immune therapy in PCa patients, the Tumor Immune Dysfunction and Exclusion (TIDE) [23] was applied. The chemotherapy response for three common drugs of each PCa patient in the TCGA and MSKCC datasets were calculated based on the Genomics of Drug Sensitivity in Cancer (GDSC, https://www.cancerrxgene.org) by performed ‘pRRophetic’ R package [24].

### 2.6. Cell culture, TFRC stably over-expressed cell lines and reagents

The HEK293T, and prostate cancer cells C4-2, PC-3, LNCaP, 22RV1 were purchased from American Type Culture Collection (ATCC, Manassas, VA). HEK293T was cultivated in DMEM while prostate cancer cells in RPMI 1640. All cells were cultured in the same humidified atmosphere (37 °C with 5% CO2). 5-Azacytidine (Sigma, #A2385) was used for the treatment of 22RV1 and LNCaP cells. The TFRC overexpressed plasmid was designed and synthesized by Genomeditech (Shanghai, China). Next, the TFRC overexpressed plasmids, psPAX2 and pMD2.G were mixed and added into HEK293T cells to package lentivirus. After 48h incubation, lentivirus supernatants were obtained, filtered and used for infecting cells.

### 2.7. Western blot analysis

After washed 3 times with cold PBS buffer, cells were lysed using RIPA buffer and kept on ice. We loaded the same amounts (40 μg) of different protein samples onto the SDS-PAGE gel and then transferred the protein to PVDF membranes. The membranes were blocked in skim milk (5%) for 1 h and incubated at 4 °C with primary antibodies overnight. The next day, we incubated the PVDF membranes with HRP-conjugated secondary antibodies (mouse or rabbit) for 1 h at room temperature. After 3 washes in TBST, the blots were visualized based on an ECL system (Thermo Fisher Scientific). Following were the primary antibodies used in this study: TFRC (ABclonal, #A5865), β-Actin (sc-517582).

### 2.8. Cell proliferation assay

MTT assay: After infected with TFRC overexpressing lentivirus, cells were seeded 1×10^3^ per well in 96-well plates. The cell viability was tested at 0, 24, 48, 72, 96 h. MTT reagent was added to each well and then the plate was incubated at 37 °C for 2 h. After the incubation, we removed the medium and added dissolved the formazan crystals in DMSO. The absorbance of each well was measured at 490 nm.

Colony formation: 22RV1 and LNCaP cells were seeded (1×10^3^ per well) in 6-well plates and allowed to grow for 12 days. Next, the culture medium was removed, and the cells were washed in cold PBS for 3 times. Then we fixed the cells using 95% ethanol for 15 min. After that, the cells were stained in 1% crystal violet for 20 mins. The colonies were counted and photographed using a microscope.

### 2.9. Cell invasion assay

After overexpressing TFRC, we used transwell chambers (8 μm pore size, Corning, MA, USA) to detect the migration and invasion of 22RV1 and LNCaP. The upper chamber was pre-coated with Matrigel (Corning, USA) for invasion and not for migration. 10×10^4^ cells with serum-free media were seeded in the upper chamber, while the lower chamber was added 600 μL 10% FBS culture media. After 12 h (migration) or 24 h (invasion), we used a cotton swab to remove the cells left in the upper surface of the chambers and fixed the cells that on the lower surface of the filters in 100% methanol for 15 min. Then, we stained them in 0.1% crystal violet solution for 20 min. The cell numbers were counted and averaged across 5 randomly chosen fields using a microscope.

### 2.10. Statistics

All data analyses in this study were performed in the R platform (v.4.0.2, https://cran.r-project.org/) or Graphpad 8.0. The comparison of mRNA expression between PCa tissues and adjacent nontumorous samples, CNV, TMB, mRNAsi, clonal score, chemotherapy response and immunotherapy response across different risk groups were calculated with Wilcoxon rank-sum test. The RFS of different groups was measured using Kaplan-Meier method with log-rank test. The DEGs between different risk groups (high and low) were utilized “limma” R package [25]. The MTT, invasion and migration results were tested via Student’s t test. *P* < 0.05 was considered as statistically significant.

## 3. Results

### 3.1. Expressions and correlations of FRGs in the TCGA cohort

Most of FRGs (32/40, 80%) (**Figure1A&1C**), (24/40,60%) (**Figure 1B&1E**) were abnormally expressed in tumor samples compared with adjacent nontumorous tissues, and paired nontumorous tissues. The interaction network among these 40 genes demonstrated that TP53, GCLC, GCLM, GPX4 and other genes have more connections (**Figure 1D**). Moreover, the correlations among these genes were very high (**Figure 1F**, for example, ACSL4 and NFE2L2, HMGCR and NCOA4, HSPB1 and CRYAB).

**Figure 1.**
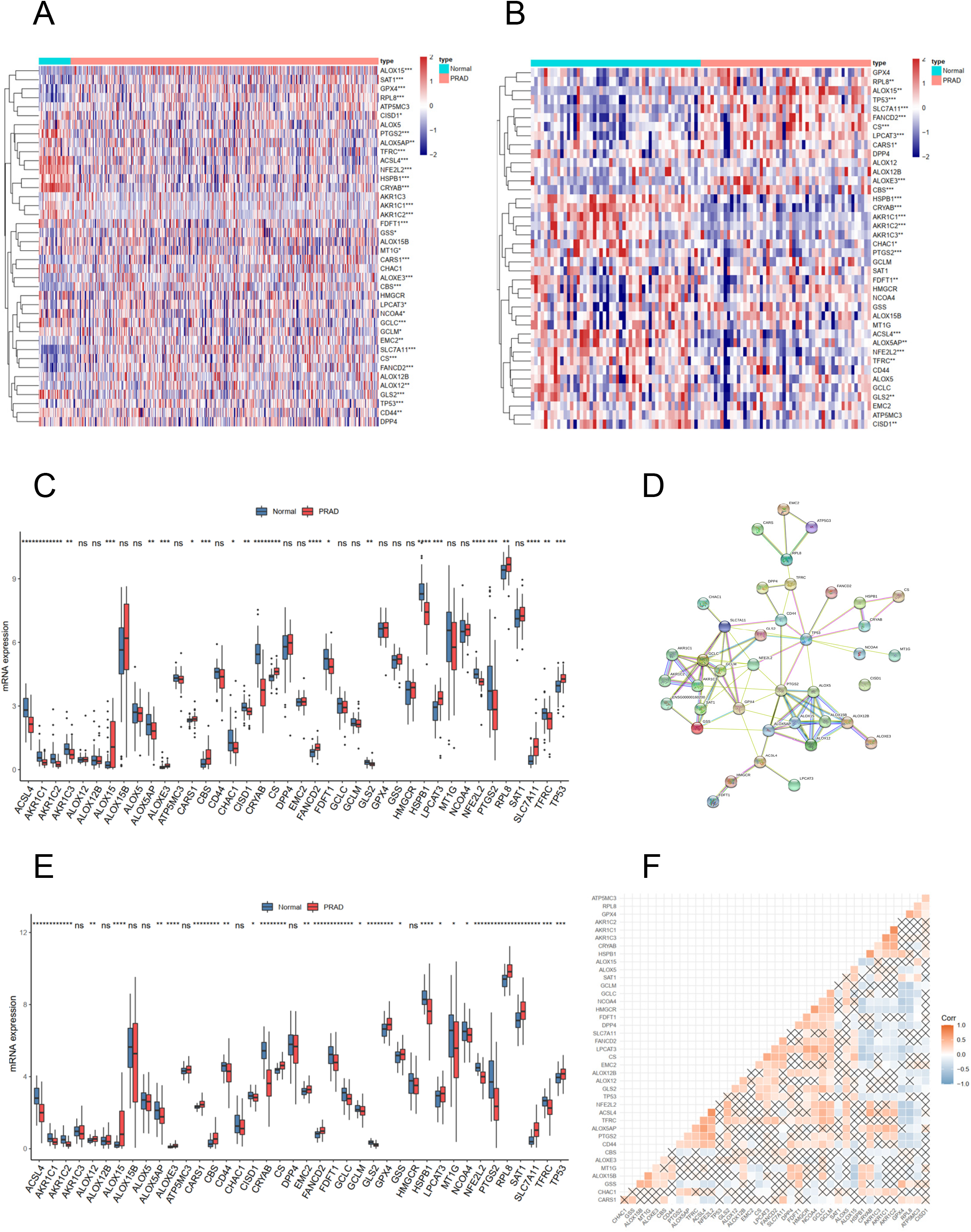
The landscape of ferroptosis-related genes (FRG) in prostate cancer. (A) Heatmap of 40 FRGs between 499 prostate cancer and normal tissues. (B) Heatmap of 40 FRGs between 52 tumor and adjacent normal pairs. (C) mRNA expression levels FRGs levels in total prostate cancer and normal tissues. (D) The PPI network acquired from the STRING database among the FRGs. (E) mRNA expression levels FRGs levels in prostate cancer and paired tissues. (F) The correlation among FRG in prostate cancer. * P < 0.05, ** P < 0.01, *** P < 0.001, ns, not significant.

### 3.2. Clustering, construction and validation of FRGs prognostic model

According to the mRNA expression levels of the 40 FRGs, the TCGA cohort could be divided into two cluster (**Figure 2A**). Moreover, patients in cluster1 had significantly poor RFS than those in cluster2 (**Figure 2B**), which suggested that FRGs could be related with the difference with PCa patients. Therefore, the explorations of the prognostic FRGs were very necessary. Firstly, the univariate Cox regression method was applied to select the prognostic-associated FRGs, which indicated that 17 genes were correlated with PCa RFS. In order to minimize the risk of overfitting, then LASSO method was utilized to choose the hub genes (**Figure 2C**). To further identify the FRGs with the greatest prognostic value, we conducted multiple stepwise Cox regression and chose seven hub FRGs to construct the prognostic model among PCa patients (**Figure 2D**). Moreover, the seven hub genes had more than 1% genetic alterations, for example TFRC (4%) and ALOXE3 (4%) (**Figure 2E**). The risk scores of each PCa patient was measured based on the format: Risk score = (0.213*ExpAKR1C3) + (0.224*ExpALOXE3) + (0.183*ExpATP5MC3) + (0.182*ExpCARS1) + (−0.346*ExpMT1G) + (−0.193*ExpPTGS2) + (0.299*ExpTFRC). Then according to the median value of risk score, we divided the PCa patients of TCGA cohorts into two groups (high and low risk).

**Figure 2.**
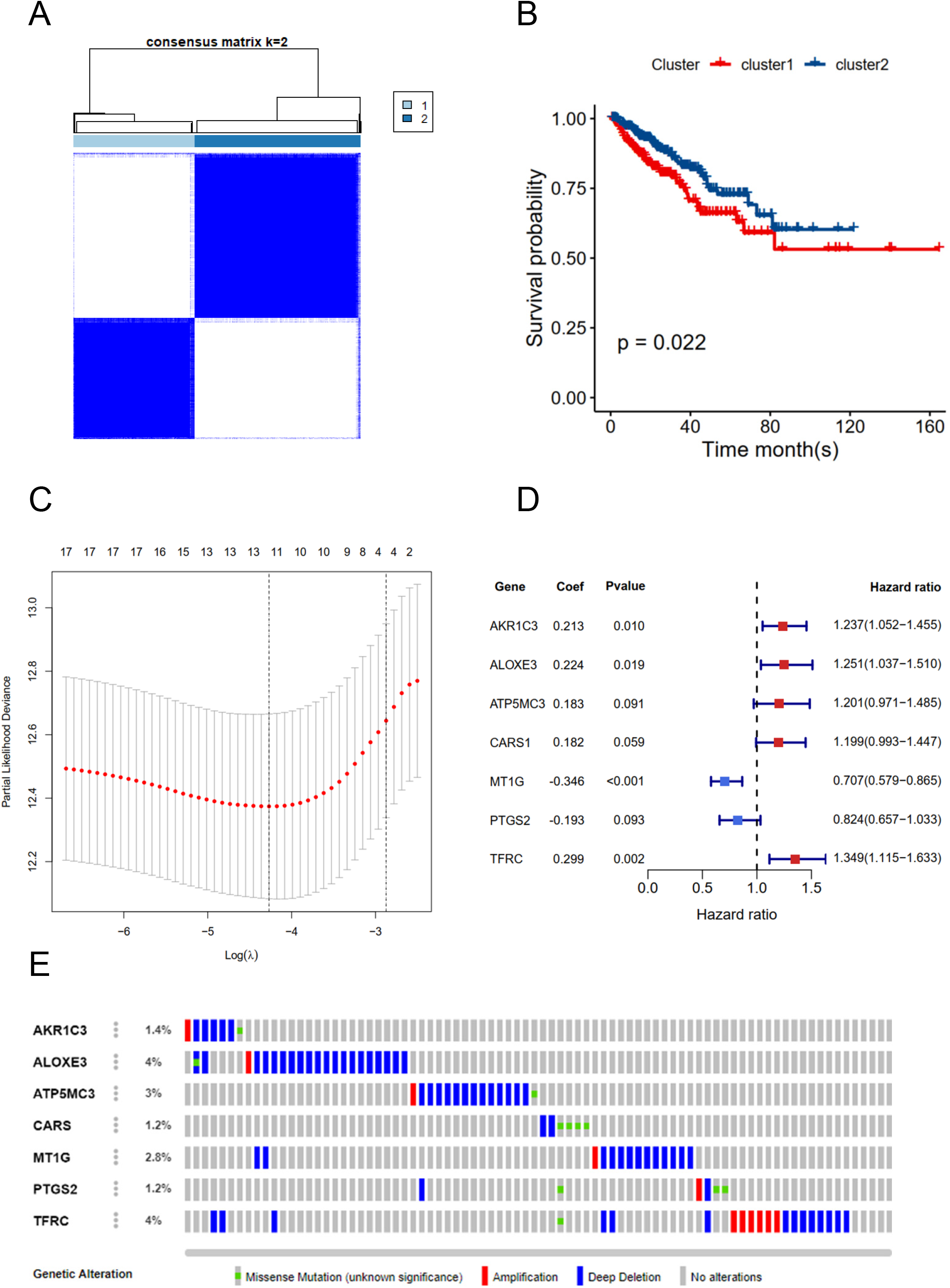
RFS of PRAD patients in cluster 1/2 subgroups and risk signature with 7 FRGs. (A) Consensus Clustering matrix for k =2. (B) Kaplan-Meier curves of DFS. (C) The Cross-Validation fit curve calculated by lasso regression method. (D) The coefficients of seven hub FRGs estimated by multivariate Cox regression. (E) Genetic alterations of seven hub FRGs.

The K-M plot demonstrated that the group with high risk had unfavorable RFS than the low risk group (P < 0.0001, **Figure 3A**). Moreover, the ROC area (AUC) under the curve in 1-year, 3-year and 5-year were 0.741, 0.729 and 0.736 (**Figure 3B**). The RFS status of PRAD, heatmap and barplot of these seven genes were showed in **Figure 3C**, **Figure 3D&3E**. Furthermore, the results of univariate and multivariate Cox regression analysis demonstrated that the risk score was an independent factor for PCa patients in TCGA cohort (HR = 1.11, 95% CI:1.08-1.15, p value < 0.001, **Table 2**). In order to verify the stability of this risk model, the MSKCC cohort was included to test the predictive value. The results were consistent with the effect of TCGA dataset (HR = 1.76, 95% CI: 1.43-2.17, p value <0.001, **Figure 4A-E and Table 2**).

**Figure 3.**
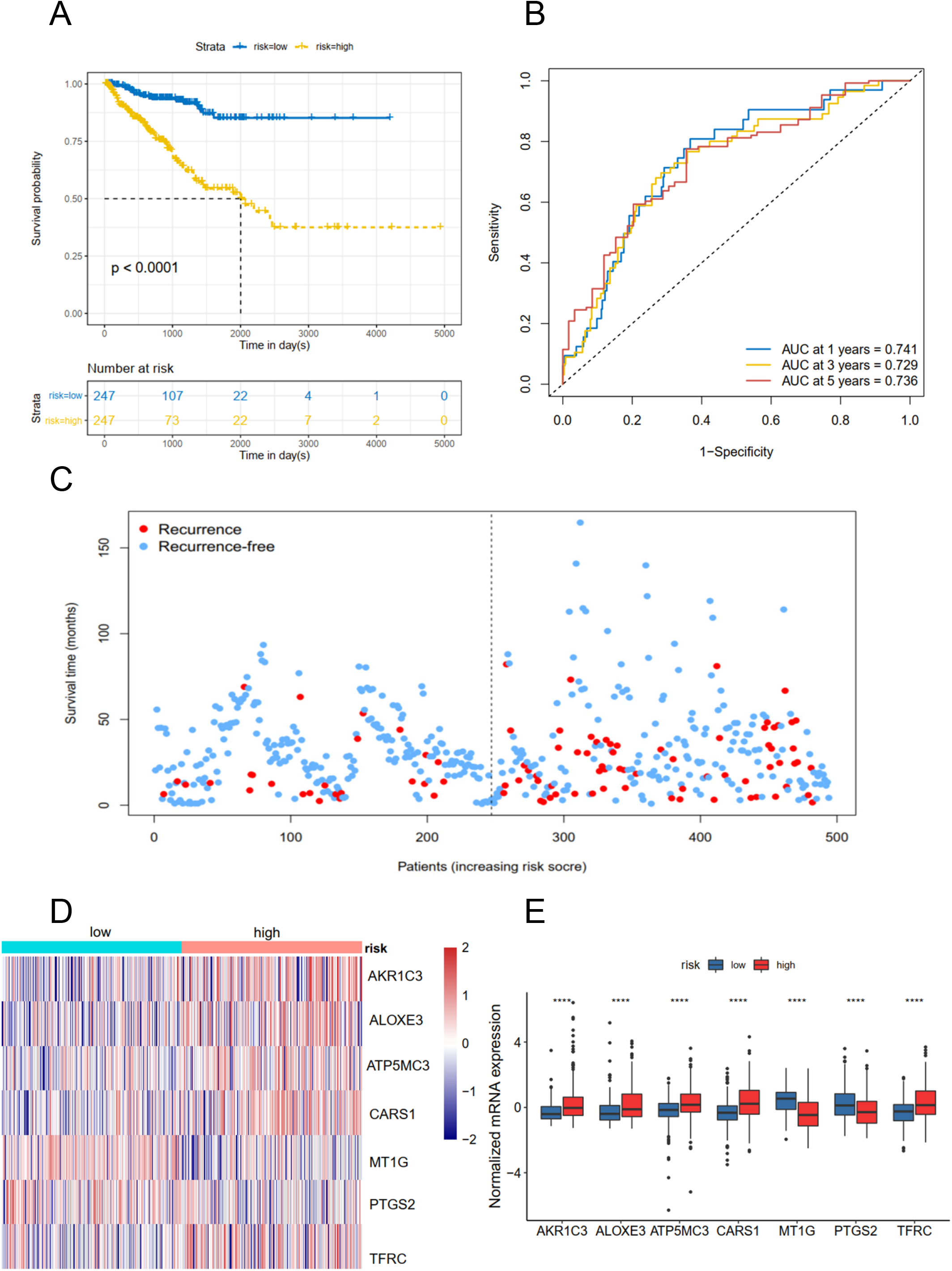
Risk score based on seven hub FRGs in TCGA PRAD cohort. (A) Survival analysis according to risk score. (B) ROC analysis. (C) Survival status of patients. (DE) Heatmap and barplot of the seven hub genes between high and low risk group.

**Figure 4.**
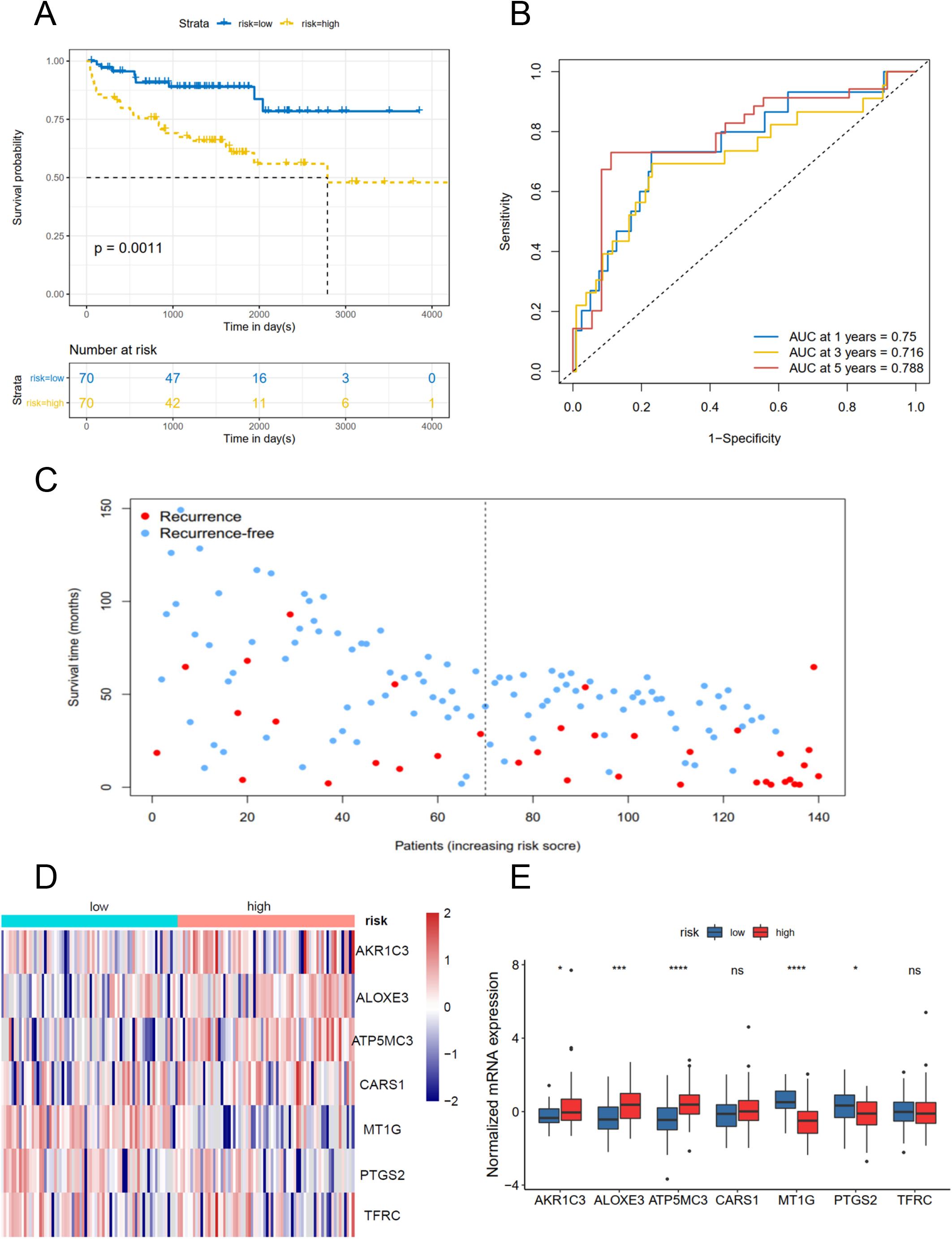
Risk score based on seven hub FRGs in MSKCC PRAD cohort. (A) Survival analysis according to risk score. (B) ROC analysis. (C) Survival status of patients. (D-E) Heatmap and barplot of the seven hub genes between high and low risk group.

### 3.3. Functional analyses in TCGA and MSKCC cohorts

To further clarify the biological functions and pathways of FRGs, the DEGs between the high risk and low risk group were analyzed in the TCGA-PCa cohort, where 664 genes reached the set (adjust P < 0.05, |log2FC| > 1), including 419 upregulated and 245 downregulated genes (**Figure 5A**). Then GO and KEGG enrichment were performed using ClusterProfiler R package. Interestingly, most of the GO terms were enriched in immune-related functions (**Figure 5B**), such as humoral immune response, regulation of humoral immune response, B cell mediated immunity, immunoglobulin complex, antigen binding and growth factor activity. While the KEGG terms were closely associated with several metabolism process (**Figure 5C**), such as ascorbate and aldarate metabolism, steroid hormone biosynthesis, especially drug metabolism and metabolism of xenobiotics by cytochrome P450, which were essential pathways for many drugs fully utilization. Considering the risk score was strongly associated with the immune response, then the 28 immune infiltration status were calculated using the ssGSEA method. The activated CD4 T cell, CD 56 dim natural killer cell, mast cell, memory B cell, neutrophil, regulatory T cell and Type 17 T helper cell were significantly distinct across low- and high-risk groups in the TCGA dataset (P <0.05, **Figure 5D**). While the CD 56 dim natural killer cell, central memory CD8 T cell, natural killer cell, natural killer T cell and Type 17 T helper cell were significantly different between groups with low or high risk in MSKCC dataset (P <0.05, **Figure 5D**). And risk scores were associated with regulatory T cell, type 17 T helper cell, CD 56 bright natural killer cell and neutrophil (**Figure 5E&5G**), which strongly prompted that FRGs may regulate the progress of PCa via the immune process pathway.

**Figure 5.**
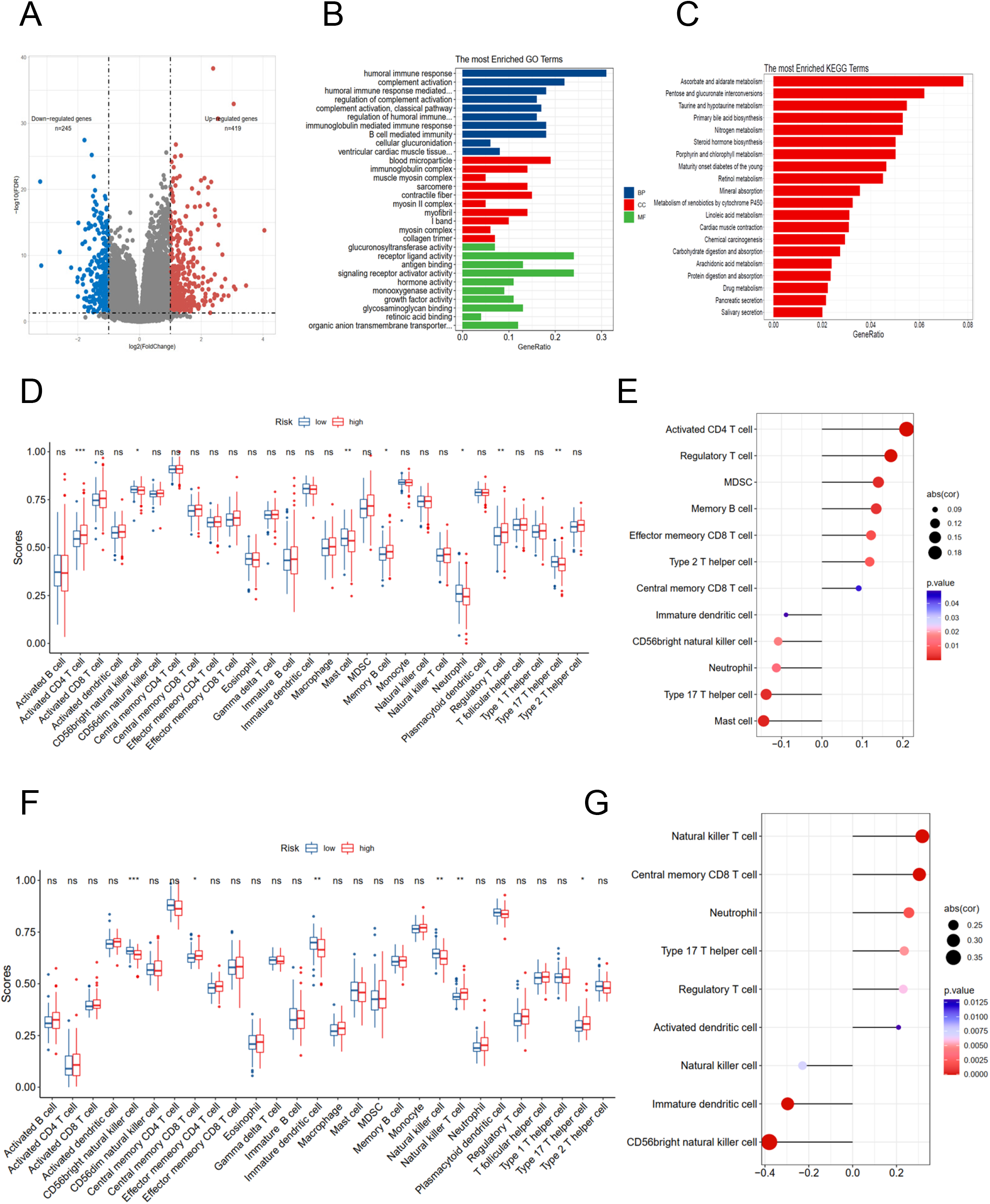
Potential biological pathways affected by FRGs. (A) The different expression genes (DEGs) between the high-risk and low-risk groups. (B) The Gene Ontology (GO) enrichment of DEGs. (C) The KEGG enrichment of high and low risk groups. Comparison and correlations of the ssGSEA scores between different risk groups and risk scores in TCGA (D-E) and MSKCC(F-G) dataset.

### 3.4. The distinctions of gene TMB, CNV, cancer stemness index and sensitivity to immuno-/chemotherapy among the groups with high- and low-risk

According to the GO and KEGG enrichment results, the risk score was closely associated with the immune process and drug metabolism pathways. Then in order to detect whether the risk score was correlated with the immune response and chemotherapy, the CNV, TMB, cancer stemness index and clonal score were included. As for the CNV status in TCGA cohort, the high risk group both had high amplication (P_amp_ = 3.3e-11, P_amp_ = 4.4e-14) and deletion (P_del_ = 2.3e-09, P_del_ = 6.7e-13) in arm and focal levels (**Figure 6A-B**). Moreover, the high risk group had high TMB (P = 1.1e-11 **Figure 6C**), neoantigens burden (P = 1.1e-05, **Figure 6D**), mRNAsi (P < 0.001, **Figure 6E**) and clonal score (P < 0.001, **Figure 6F**) in TCGA dataset. Notably, TMB and neoantigens were potential biomarkers for immunotherapy response [26]. Subsequently, the immunotherapy response and predictive responses for three common chemotherapy drugs, cisplatin, docetaxel and bicalutamide were analyzed based on the TIDE and GDSC database. The results revealed that TIDE score was increased in high risk group (P = 0.032, TCGA dataset **Figure 6G**), though there is no significantly different in MSKCC dataset (P = 0.81, **Figure 6I**). The group with high risk both had high estimated IC50 for Bicalutamide (P = 8.5e-11 TCGA; P = 1.5e-05 MSKCC), and low estimated IC50 for Docetaxel (P = 0.025 TCGA; P = 0.076 MSKCC), but not for Cisplatin (**Figure 6H&6J**).

**Figure 6.**
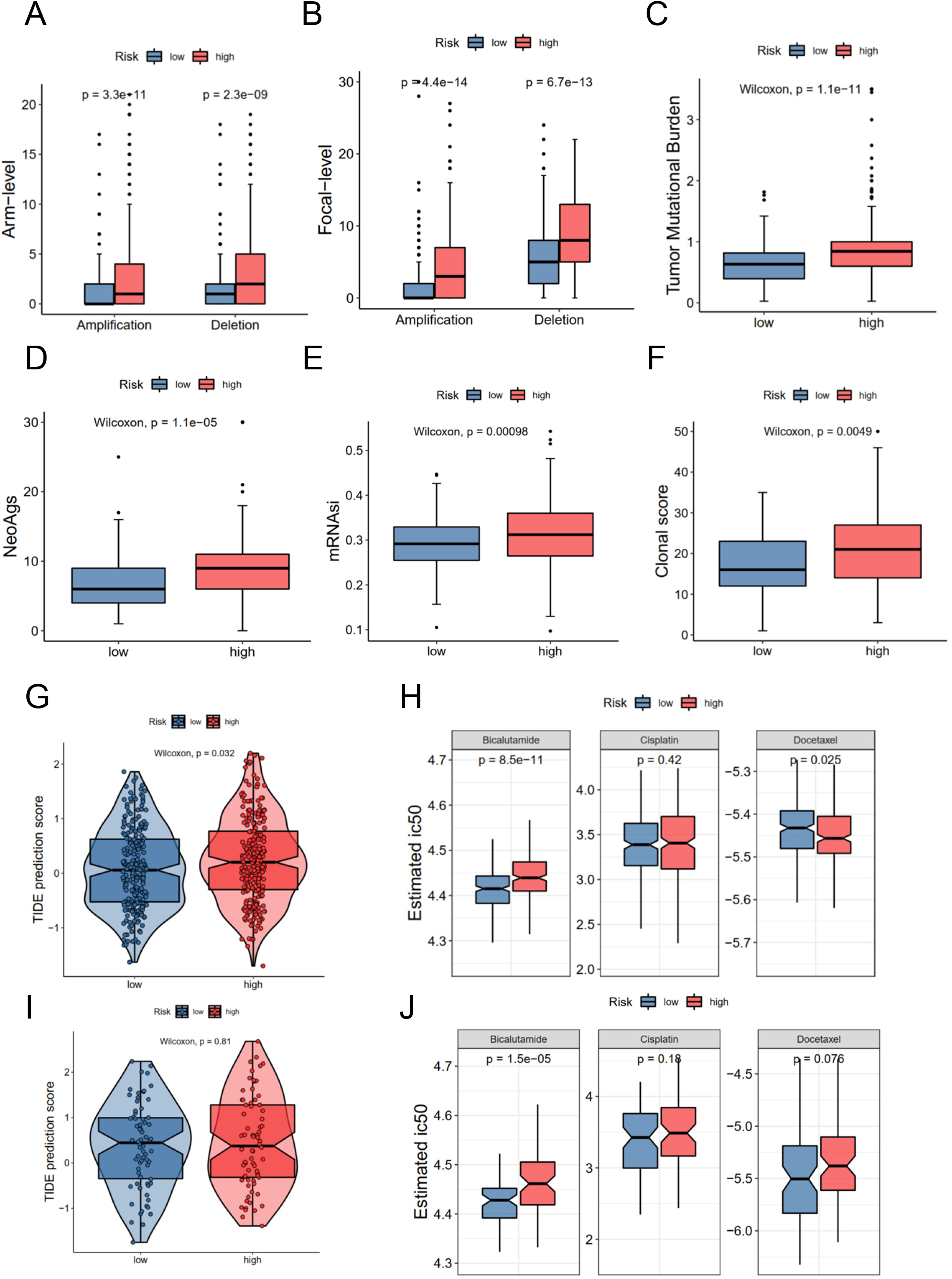
The correlations of different risk group with copy number alterations, tumor mutation burden, neo-antigens, tumor stemness, clonal status and immuno-/chemotherapy response. (A) Arm-level copy number amplification and deletion. (B) Focal-level copy number amplification and deletion; (C) Tumor mutant burden difference. (D) Neo-antigens. (E) Tumor stemness difference represented by the mRNAsi. (F) Clonal status. (G-H) Immunotherapy response based on TIDE website and estimated IC50 indicates the efficiency of chemotherapy in TCGA. (I-J) Immunotherapy response and estimated IC50 in MSKCC dataset.

### 3.5. Overexpressing TFRC facilitates proliferation, migration and invasion in PCa cell lines

Considering the risk model was strongly associated with the RFS of PCa, then the hub genes may have more effect on the biological functions. Therefore, we chose the high genetic alteration gene TFRC to confirm our hypothesis. Firstly, we measured the baseline protein levels of TFRC in PCa cells PC-3, LNCaP, 22RV1 and C4-2 by western blots. The expression of TFRC was relatively lower in LNCaP and 22RV1 cells (**Figure 7A**). Thus, we used these two cell lines to overexpress TFRC in later experiments. Interestingly, the mRNA expression of TFRC decreased in PCa tissues (**Figure 1C&1E**) when compared to normal samples, but increased in high gleason score tissues and lymph nodes metastasis (**Supplementary S1A&1B**). Several literatures reported that methylated CpG sites can block transcription initiation via inhibiting the binding of transcription factors, like promoters and distal regulatory regions, then change the mRNA expression of host genes [27–29]. Then we explored the methylation levels of TFRC and the results showed that TFRC had high methylation levels in PCa tissues (**Supplementary S1C**), which were negatively associated with the mRNA expression of TFRC (**Supplementary S1D**). Therefore, we treated the PCa cells with 5-zacytidine (a widely-used DNA methylation inhibitor [30]) and found that the protein level of TFRC was increased after cells were incubated with 0.5 μM 5-zacytidine for 24 h (**Figure 7B**). Next, we overexpressed TFRC in 22RV1 and LNCaP cells (**Figure 7CD**). The effect of TFRC on PCa cells were evaluated by MTT, colony formation and transwell assays. As shown in **Figure 7E-F**, upregulation of TFRC induced proliferation in PCa cells. In parallelly, the transwell assays indicated that TFRC promotes the migrative and invasive activities of LNCaP and 22RV1 cell lines (**Figure 7G**).

**Figure 7.**
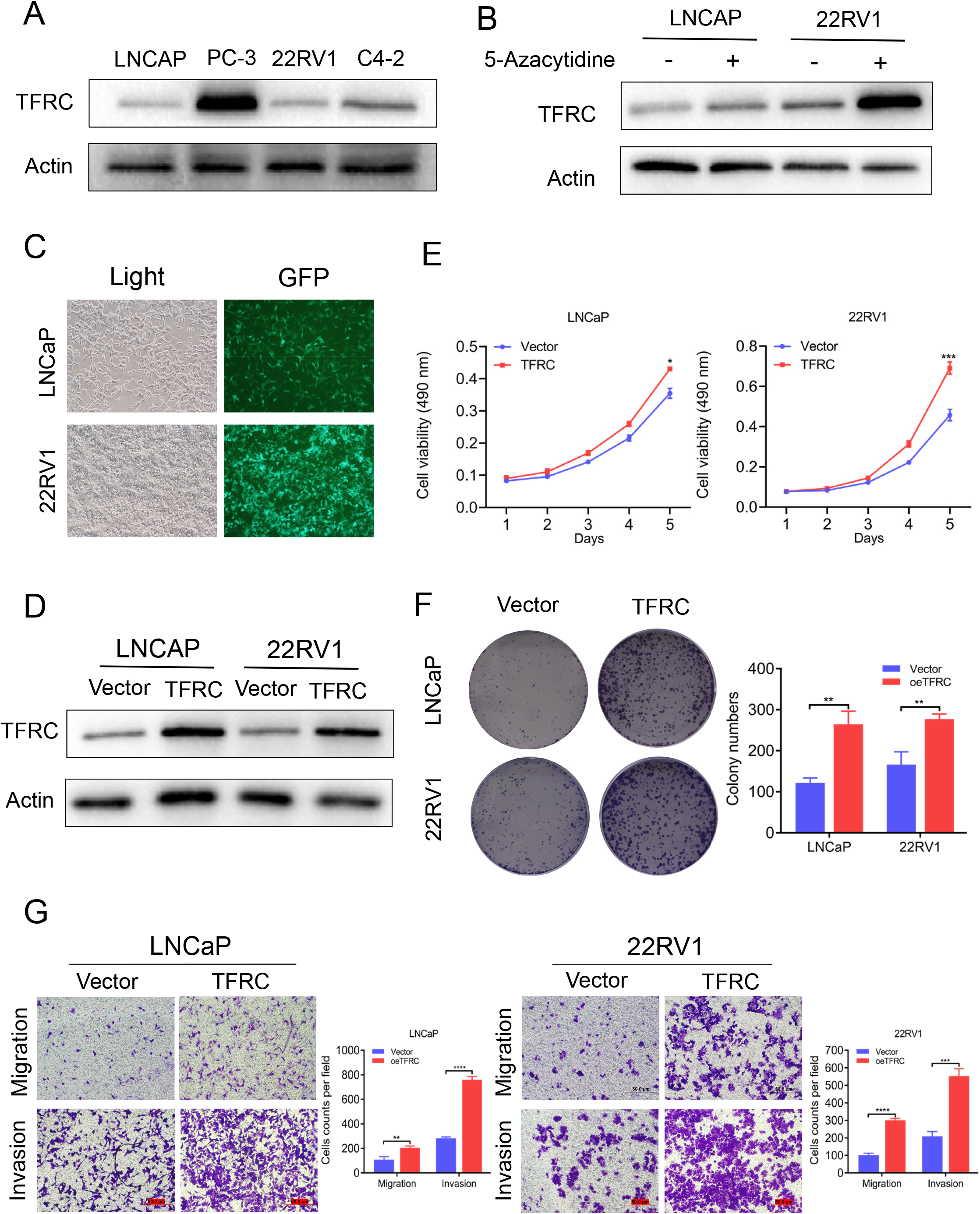
TFRC expression influenced by 5-zacytidine and overexpressing TFRC promotes proliferation, migration and invasion in PCa cells 22RV1 and LNCaP. (A) The protein levels of TFRC in PCa cells (C4-2, LNCaP, 22RV1 and PC-3) were detected by western blots. (B) 5-azacytidine inhibited DNA methylation and increased the expression of TFRC in LNCaP and 22RV1 cells. (C) and (D). The efficiency of TFRC overexpressed lentivirus were confirmed using fluorescence microscope and western blots. (E) and (F). The MTT and colony formation assays indicated that TFRC promotes cells proliferation in LNCaP and 22RV1. (G) and (H). Transwell assays suggested that TFRC promotes cells migration and invasion in LNCaP and 22RV1.

## 4. Discussions

Selective induction of cancer cell death is the most effective therapy method for tumor [31]. Several literatures reported that ferroptosis, a common selective induction cell death, plays pivotal role in the process of tumorigenesis [32, 33]. However, the systematic analysis of prostate cancer has yet to be elucidated. In this study, we collected the RNA-seq data to investigate expression variations in mRNA expression profiles of ferroptosis-related genes in prostate cancer. On the basis of unicox, lasso regression and multicox analyses, the signature of seven ferroptosis-related genes was identified.

It was widely reported that the seven hub genes, AKR1C3, ALOXE3, ATP5MC3, CARS1, MT1G, PTGS2, and TFRC were involved in the development of several diseases. AKR1C3, a crucial androgenic enzyme, could reprogram AR signaling in advanced prostate cancer [34] and implicated in the production of aromatase substrates in breast cancer [35]. ALOXE3, epidermal LOX type 3, converting fatty acid substrates through R-hydroperoxides to specific epoxyalcohol derivatives [36], involved in late epidermal differentiation [37] and ichthyosis [38]. ATP5MC3, also named ATP5G3 [39], encoding a subunit of mitochondrial ATP synthase, was associated with overall survival in clear cell renal carcinoma patients [40]. CARS1, Cysteinyl-TRNA Synthetase 1, associated with tRNA function, has been found associated with the development of inflammatory myofibroblastic tumor [41] and kidney cancer [42]. CARS1 knockdown has been proved to suppress ferroptosis induced by cysteine deprivation and promote the transsulfuration pathway [43]. MT1G, a small-molecular weight protein that has high affinity for zinc ions, could inhibit the proliferation via increasing the stability and the transcriptional activity of p53 [44]. PTGS2, also known as cyclooxygenase 2, is responsible for the prostanoid biosynthesis [45]. Several researches demonstrated that PTGS2 induces cancer stem cell like activity and promotes proliferation, angiogeniesis and metastasis of cancer cells [46–48]. As for TFRC, a cell surface receptor necessary for cellular iron [49]. Notably, except PTGS2 and MT1G, few of these genes, has ever been reported in PCa. Here in this study, these seven genes were demonstrated correlated with the RFS of PCa.

Based on the mRNA expression profiles of these seven hub genes, the ferroptosis related risk model was constructed in TCGA PCa cohort. The KM plot and ROC curve showed this risk score could easily distinguish different RFS. Moreover, the results were validated in the MSKCC external validation dataset. To further explore the detail mechanism of the FRGs, the PCa patients were distributed into two groups according to the risk score. There were 664 DEGs between these two different groups. Interestingly, these DEGs were enriched in immune related GO terms and metabolism KEGG terms (**Figure 5B-5C**). Subsequently, the immune infiltration status of 28 immune cells were calculated based on the ssGSEA method, and the results showed that several immune cells were associated with risk score (**Figure 5D-5G**), such as central memory CD8 T cell, CD56 bright natural killer cell, which suggested this risk model may have prompt for immunotherapy. Then, the immunotherapy related signature, such as CNV load, TMB, mRNAsi, clonal score, and TIDE score were included for further investigation. These results indicated that high risk group was correlated with CNV load, TMB, neoantigens, mRNAsi and clonal score (**Figure 6A-6F**), which suggested that high risk score group may have better response for immune therapy. The TIDE score in high risk group was also increased, which also suggested the better outcome for immune therapy (**Figure 6G&6I**). Besides, considering the most KEGG terms between low and high risk group were enriched in metabolism, especially in drug metabolism (**Figure 5C**), we detected the estimated IC50 for three commonly used drugs. The results showed an increasement in high risk group for Bicalutamide (**Figure 6H-J)**, which implied that this group may be resistant for Bicalutamide. In a word, the FRGs may have effects on the RFS via regulation of the drug response, however, the high risk patients may improve their situation through different immune therapy, which may provide a potential strategy in individual treatment of PCa patients. Finally, we chose the high genetic alteration rate gene TFRC for further validation. Considering the mRNA level of TFRC is low in PCa patients when compared with normal tissues, we decided to overexpress TFRC to detect the effect on PCa cells. Based on the baseline protein expression, LNCaP and 22Rv1 cell lines were selected (**Figure 7A**, **Figure 7C** and **Figure 7D**). MTT assay, clonal formation and invasion showed that TFRC overexpression could significantly increase the proliferation (**Figure 7E, 7F**) and invasion (**Figure 7G**) in PCa cells. Moreover, we also add 5-Aza to reduce the DNA methylation of TFRC, which demonstrated that the TFRC gene may have hypermethylation status and thus induce the relatively lower expression in PRAD tissues, but rising trend along with the gleason score. This suggests that TFRC may play an oncogenic role in PCa, but the detail mechanism needs to be explored further.

In summary, our study systematically analyzed the expression of FRGs and their potential prognostic ability in PCa. Then, the risk model of seven FRGs was established in TCGA and validated in MSKCC dataset. Moreover, we detected that high risk group was correlated with high CNV load, TMB, neoantiogens, mRNAsi, clonal score and immune therapy response, but with high estimated IC50 for Bicautamide. Finally, the impacts on proliferation and invasion when overexpressing TFRC in PCa cell lines were tested. Our research has several limitations. First, the risk model in our study was established and validated both based on public databases. However, to confirm its clinical significance, more prospective real-world data are needed. Second, we only initially explore the effect for overexpression of TFRC on biological functions, the specific mechanism needs to be discussed further *in vivo* and *vitro.*

## 5. Conclusions

In conclusion, a ferroptosis-related risk model was established, which were strongly associated with aberrantly CNV load, TMB, mRNAsi, neoantigens, clonal score, immune-/chemo-responses. Simultaneously, TFRC is validated to be an oncogenic factor in prostate cell lines. Our research provides new insights of the personalised therapy for PCa patients.

## Author statement

**Yunfei Xu**, **Lei Gao**: Conceptualization; Supervision; Methodology; Writing -review & editing. **Huan Liu**, **Lei Gao**: Data curation; Formal analysis; Writing-Original draft. **Jie Li**, **Tingshuai Zha**i and **Tiancheng Xie**: Validation; Visualization; Investigation.

## Data accessibility

All data about TCGA and MSKCC dataset are publicly available, and appear in the submitted article.

## Acknowledgments

This study was sponsored by the National Natural Science Foundation of China (NO. 81671446, NO.81971371).

---

**Supplementary Figure S1.** The associations between TFRC mRNA expression levels and gleason score (A), nodal metastasis (B) and methylation levels (C-D) in TCGA PCa patients.

