## Supplementary figures and images for "Identification and validation of a ferroptosis-related genes based prognostic signature for prostate cancer"

### Supplemental figure1

#### Supplementary Figure S1

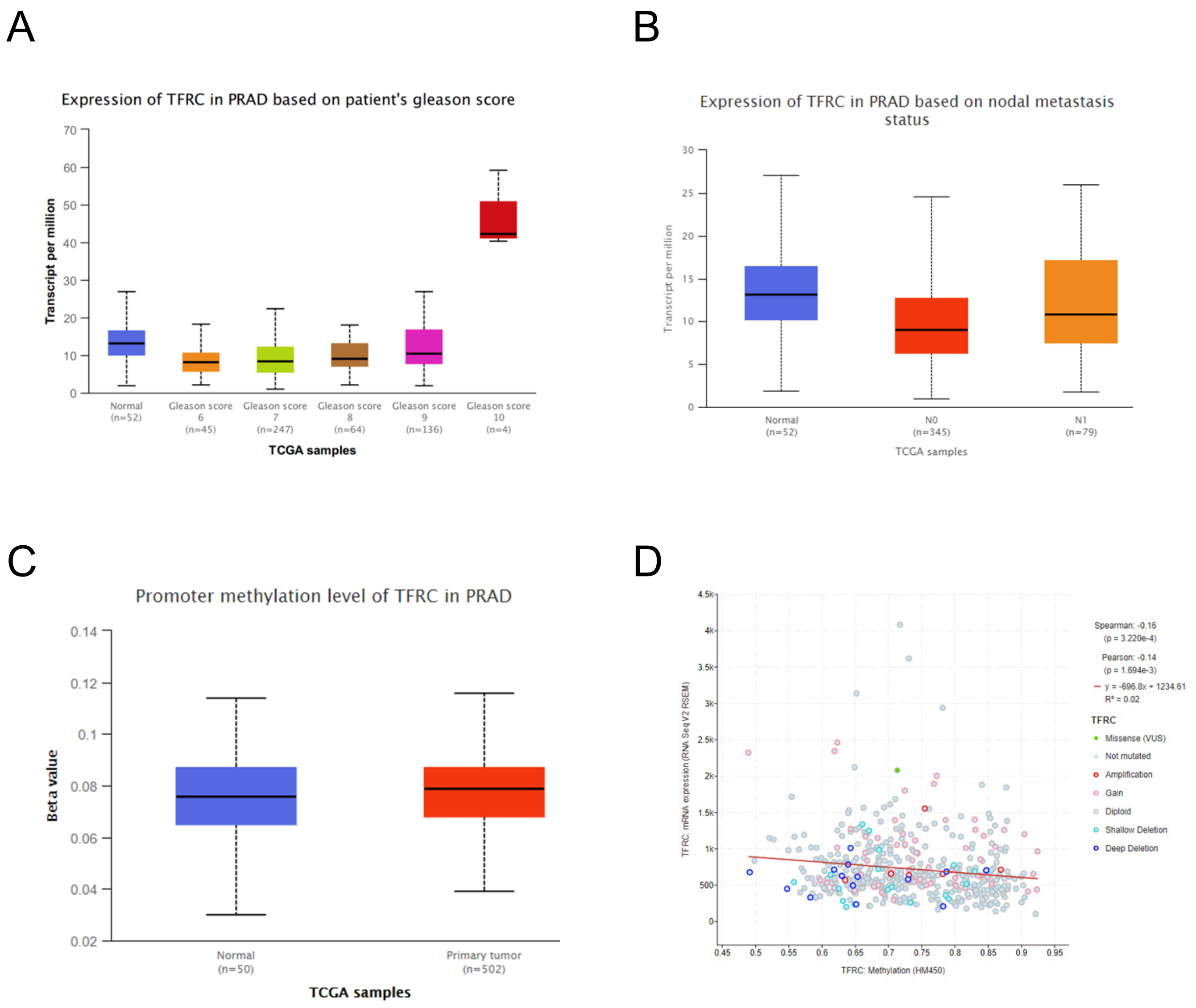
